# A Resident State Allows Influenza Polymerase to Smoothly Switch between Transcription and Replication Cycles

**DOI:** 10.1101/2021.11.30.470684

**Authors:** Huanhuan Li, Yixi Wu, Minke Li, Lu Guo, Quan Wang, Zhaohua Lai, Jihua Zhang, Xing Zhang, Lixin Zhu, Ping Lan, Zihe Rao, Yingfang Liu, Huanhuan Liang

## Abstract

Influenza polymerase (FluPol) transcripts viral mRNA and switches to replicate viral genome after transcription. However, it remains unknown how FluPol switches between transcription and replication cycles, especially when considering that the structural basis of these two functions is fundamentally different. Here, we proposed a mechanism that FluPol achieves the functional switching between these two cycles through an unreported intermediate conformation, termed as resident state. We obtained a resident state structure of H5N1 FluPol at 3.7 Å using cryo-EM, which is characterized by a blocked Cap-binding domain and a contracted core region, distinct from the structures of either transcription or replication states. Structural analysis results suggest that the resident state structure is feasible to smoothly transit into structures of both transcription and replication states. Furthermore, we show that formation of the resident state is required for both transcription and replication activities of FluPol. Together, the transcription and replication cycles of FluPol are connected via a resident state.

## Introduction

In prokaryotic and eukaryotic cells, the replication and transcription of genome are respectively executed by DNA polymerase and RNA polymerase. However, such distinct functions are often performed by a self-encoded viral polymerase in viruses including lethal viruses that cause global public health problem, such as influenza, Ebola, measles, rabies, RSV, etc. The influenza virus was studied as a representative model for negative-stranded RNA viruses. The influenza viral genome is packaged into 8 ribonucleoproteins (RNPs) by multiple copies of viral nucleoproteins (NPs) together with a heterotrimeric RNA-dependent RNA polymerase (RdRp, also termed as FluPol) (*1-3*). The FluPol composed of subunits PA, PB1 and PB2 is responsible for both viral transcription and replication (*4, 5*). It was reported that FluPol preferentially synthesizes viral mRNA for production of viral proteins including FluPol and NP at early stage of infection (*6, 7*). Once newly synthesized FluPol have accumulated, transcription will gradually switch into genome replication mode. Recently, the emerging structural studies on FluPol advanced the understanding of its respective transcription and replication mechanisms (*5, 8-14*).

The structure of FluPol is divided into two parts. The lower part is the catalytic core region for RNA synthesis including the C-terminal domain of PA (PA-C), PB1, and the N-terminal of PB2 (PB2-N), while the upper part is the regulatory region, including the N-terminal domain of PA (PA-N) and the C-terminal domains of PB2 (PB2-C), which consists Lid, Mid, Cap-binding, the domain harboring residue K627 (627 domain) and NLS domains (*15-17*). Previous structural studies showed that FluPol displays highly structural flexibility in transcription and replication cycles (*9, 18*). FluPol displays multiple conformations for its multiple functions by rotating, translating and arrangement of PA-N and PB2-C domains, while the core region is considered to be relatively stable (*15, 18, 19*).

The structural flexibility of FluPol has been illustrated at each step of its transcription state (T-state), including pre-initiation, initiation, elongation, termination and recycling phases (*12, 14*). In the pre-initiation structures, the 627 domain of PB2-C is tightly bound to the core region. The 627 domain and PA-C provide a series of continuous binding sites for the C-terminus of Pol II (Pol II CTD) (*10, 11, 20*). The collaboration between PA-N and Cap-binding domain is important for “cap-snatching” in T-state. The Cap-binding domain is exposed and favors binding of the 5′-cap of host cellular mRNA provided by Pol II (*5, 21*). The endonuclease active site of PA-N locates opposite to the Cap-binding site, the distance between which is approx. 50 Å, sufficient for PA-N to produce an oligo primer of approx. 10-15 nt length (*22-24*). To initiate transcription, the Cap-binding domain rotates ∼70° towards the catalytic cavity of PB1, which guides the primer stretch into the catalytic center. In addition, it was reported that the free 424-loop leads the capped primer to the catalytic channel of polymerase during transcription (*5, 20*). Following the synthesis and elongation of the newly synthesized mRNA, several functional motifs of PB1, including motifs pre-A/F, B, C and E are pushed out of the cavity by template-mRNA duplex. During termination, FluPol stutters on an oligo(U) motif starting at position U17 from the 5′ terminus and generate the poly(A) tail (*14*). To fulfil a complete transcription cycle, FluPol would be recycled through a structural reconfiguration process, which remains unclear (*14*).

Unlike CapRNA-primed initiation of transcription, FluPol initiates replication by *de novo* synthesis of the dinucleotide pppApG (*25*). Previous studies have shown that at least two FluPols, one replicating FluPol (replicase) and one recruited FluPol (encapsidating FluPol) are required to initiate the replication process (*8, 13*). ANP32A, a host factor, was identified to be required in FluPol replication (*26, 27*). In the structure of ANP32A-mediated FluPol dimer (*13*), the structure of replicating FluPol is distinct from the structures shown in T-state. The 627 domain of replicating FluPol locates more distant from the core region. This conformational change is detrimental for the loading of Pol II CTD, but favorable for the recruitment of ANP32A (*18*). More importantly, the Mid domain of replicating FluPol partially inserts into the active pocket of Cap-binding domain, thus blocking the binding of CapRNA to this domain (*9, 16*). Simultaneously, PA-N rotates ∼90° *in situ*, facing opposite to the Cap-binding domain. Consequently, FluPol in replication state (R-state) is unable to perform “cap-snatching” (*16*). Although it is still unclear that how FluPol make structural changes relate to replication elongation and termination, it is explicit that FluPol adopts completely different conformations to initiate transcription and replication cycles (*12-14*).

Several studies have proposed an efficient switching mechanism between transcription and replication states which is essential during influenza infection (*6, 7, 28, 29*). However, it remains unknown how the fundamental conformational changes of FluPol between T-state and R-state occurs. In particular, previous studies have shown that FluPol itself is able to fulfill transcription and replication *in vitro* (*4, 14, 30, 31*), suggesting that it could complete the switching independent on other host or viral factors. The structural determination of FluPol makes this question more prominent. Since the T-state differs fundamentally in its structure from the R-state, severely steric clashes between Cap-binding and 627 domains would ensue when FluPol directly transits from T-state structure to R-state structure (*9, 13*). Therefore, we speculate that there is one or more intermediate conformations to connect the transcription and replication cycles. The structure of the intermediate conformation needs to meet two criteria as follows. Firstly, it is the unknown structure which is distinct from either the T-state structure or the R-state structure. In addition, after a small conformational change, it can be easily converted into either a T-state structure or a R-state structure. Here, we present a H5N1 avian influenza RdRp (FluPol_H5N1_) structures that meets the above criteria, and verified its function using polymerase activity assays. Our results suggest that the intermediate conformation, which was termed as resident state, mediates the conformational switch of transcription and replication cycles.

## Results

### The structure of FluPol_H5N1_ represents a resident state

To capture the intermediate conformation of FluPol, we reconstructed multiple H5N1 avian influenza RdRp (FluPol_H5N1_) structures using cryo-EM 3D reconstruction. We obtained a structure consists a full-length of FluPol_H5N1_ bound to the vRNA promoter, which is different from the reported structure. The structure was refined to yield an EM map at the average resolution of 3.7 Å (Fourier shell correlation at 0.143 criterion). In addition, the core region of the structure consisting of PA-C, PB1 and PB2-N was individually refined to 3.6 Å as an intermediate step to assist model building of the full-length structure (Fig. S1 and S2). The final FluPol_H5N1_ structure presents as a monomer, covering all the three subunits except for a small NLS domain (residues, PB2 680-C), probably due to its highly structural variability. The PA-N and PB2-C domains of the FluPol_H5N1_ locate at the upper part of the structure while PB2-N and PA-C clamp PB1 tightly, forming the catalytic core responsible for RNA synthesis (Fig. 1a). The 5′ terminus of the vRNA promoter shows the typical hook structure bound in a pocket formed by PA-C and PB1 as previously observed (*4, 5*). However, the 3′-vRNA end (1-UCGUUU-6) points away from the PB1 cavity and rests outside the template entrance of polymerase (Fig. 1a), which was reported as resting conformation (*5*), needing to translocate into the PB1 active site to initiate transcription and replication as template (*13, 22*). Conserved residues, Arg38 and Arg46 in PB2, as well as residues in PB1 including Arg135, residues 671–676 and especially those in the PB1 β-ribbon are involved in stabilizing the 3′-vRNA end (Fig. S3).

**Figure 1.**
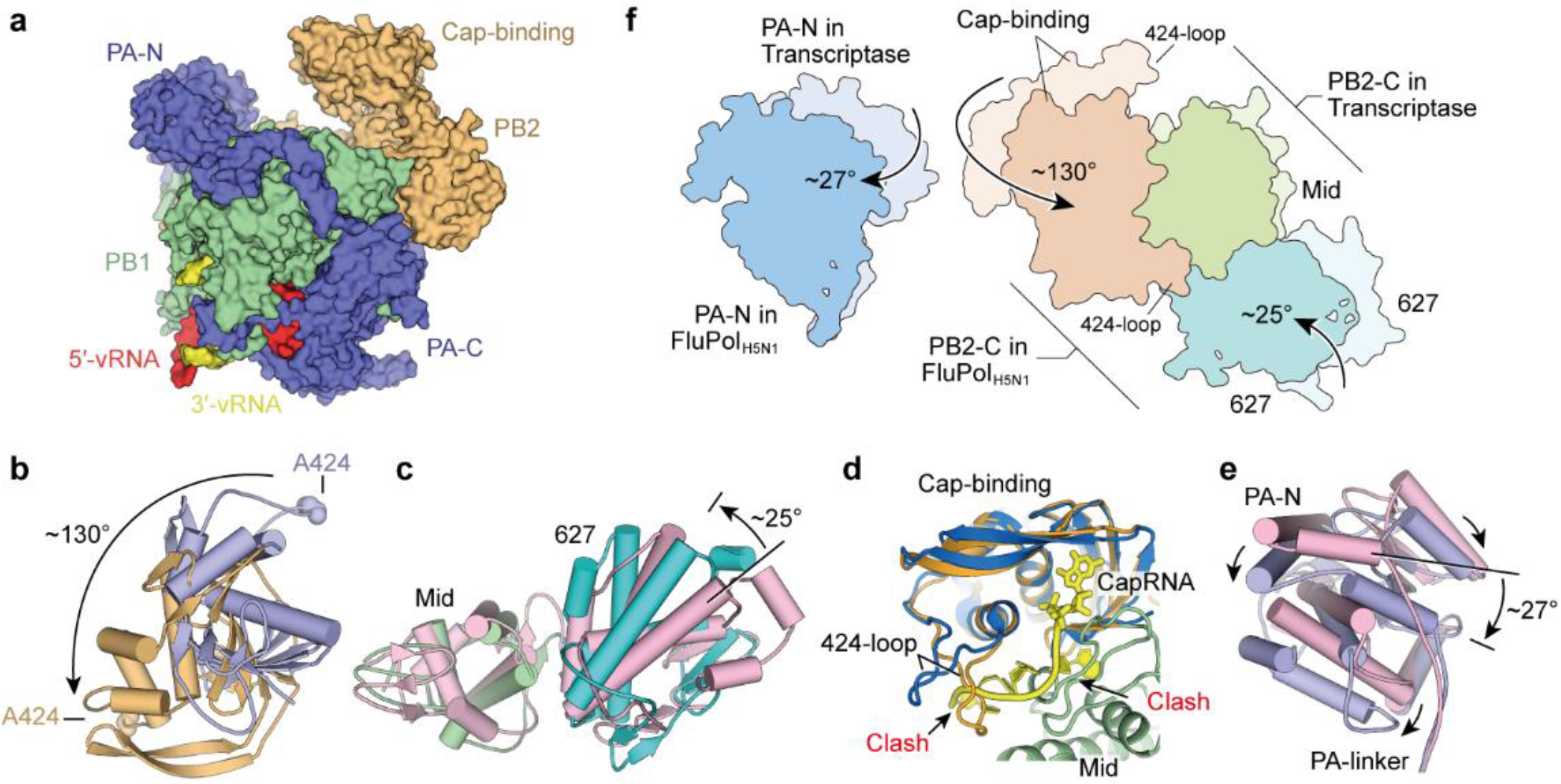
Transition from resident FluPol_H5N1_ state to transcriptase state. **a**. The cryo-EM structure of FluPol_H5N1_ complex bound with vRNA promoter. The PA, PB1 and PB2 subunits are colored in blue, green and orange, respectively. The 5′-vRNA is shown in red and 3′-vRNA is shown in yellow. **b**. Large rearrangements of Cap-binding domains between FluPol_H5N1_ and transcription pre-initiation state. The indicating residue Ala424 is shown as spheres. **c**. Rotation of 627 domains after superposition of FluPol_H5N1_ (cyan) and transcriptase (pink) structures. **d**. The 424-loop in the Cap-binding domains of FluPol_H5N1_ (orange) sterically clashes with the position of the docked 5′-CapRNA (yellow). **e**. Superposition of FluPol_H5N1_ (indicated in blue) and transcriptase FluPol (PDB: 4WSB) reveals positional and rotational changes of the PA-N domains. **f**. The conformational changes of PA-N and PB2-C domains between FluPol_H5N1_ and transcriptase are shown as models. PB2-C domains were tightened and the distance between PA-N and PB2 cap-binding domain was widened (also see Figure S4). Domains in FluPol_H5N1_ are indicated in darker color, while domains in transcriptase are shown in lighter color.

The structure of FluPol_H5N1_ is distinct from the reported T-state structures. The major difference of the FluPol_H5N1_ structure to that in T-state is a contraction of PB2-C caused by the interactions between its internal domains (Fig. 1f). Compared with those domains observed in the transcription pre-initiation states (*4*), 627 domain and Cap-binding domain in FluPol_H5N1_ rotate ∼25° and ∼130° toward its Mid domain (Fig. 1b, 1c, 1f), respectively, resulting in a highly compact PB2-C domain. Consequently, the cap binding pocket in FluPol_H5N1_ is blocked by Mid domain. As shown in Fig. 1d, when docking the 5′-CapRNA into the cap binding pocket of FluPol_H5N1_, we found that the RNA clashed with the 424-loop of the Cap-binding domain. In addition, the 424-loop is locked in the FluPol_H5N1_ structure by interacting with 627 domain (Fig. 3a). Furthermore, the PA-N domain rotates ∼27° away from the Cap-binding domain (Fig. 1e, 1f). As a result, the distance between these two domains is increased to ∼67 Å in the FluPol_H5N1_ structure, far more than those in the transcription pre-initiation state and transcription initiation state, the ∼55 Å and ∼46 Å, respectively (*4*),(*22*) (Fig. S4). The collaboration between PA-N and Cap-binding domain for “cap-snatching” in T-state is destroyed in the FluPol_H5N1_ structure. Thus, the FluPol_H5N1_ structure represents a transcription inactive state.

The structure of FluPol_H5N1_ is also different from the reported R-state structure. The major difference of FluPol_H5N1_ structure to that in R-state is a rotation of the entire PB2-C domain (Fig. 2a). The compact structure of PB2-C domain in FluPol_H5N1_ is similar to that in the replicating FluPol (*13*), whose Cap-binding domain is partial blocked by the Mid domain. However, compared with the structures of FluPol_H5N1_, the entire PB2-C domains of replicating FluPol rotates ∼130° towards the PA-N domain, basically taking the Cap-binding domain as the rotation pivot (Fig. 2a). This rotation of approx. 130° results in a protruding of the 627 domain towards the solvent (Fig. 2f). In this protruding 627 domain, the binding sites of ANP32A are fully exposed, which is important for the assembly of replicase (*13, 18*). In contrast, the ANP32A binding sites are partially blocked by the core region of the FluPol_H5N1_ structure. In addition, the PA-N domain in replicating FluPol rotates ∼90° *in situ*, opposite to the Cap-binding domain. The rotation of PA-N expands space to accommodate the NLS domain of PB2-C (Fig. 2c, 2d, 2e). Although the structure of the NLS domain in our structure was not visible, it is presumably similar to the structure found for the R-state. Rotating the PB2-C of the FluPol_H5N1_ structure for ∼130° according to that in the replicating FluPol, we found that the presumed NLS domain would sterically clash with the PA-N domain (Fig. 2d). These results show that our FluPol_H5N1_ structure represents a replication-inactive state.

**Figure 2.**
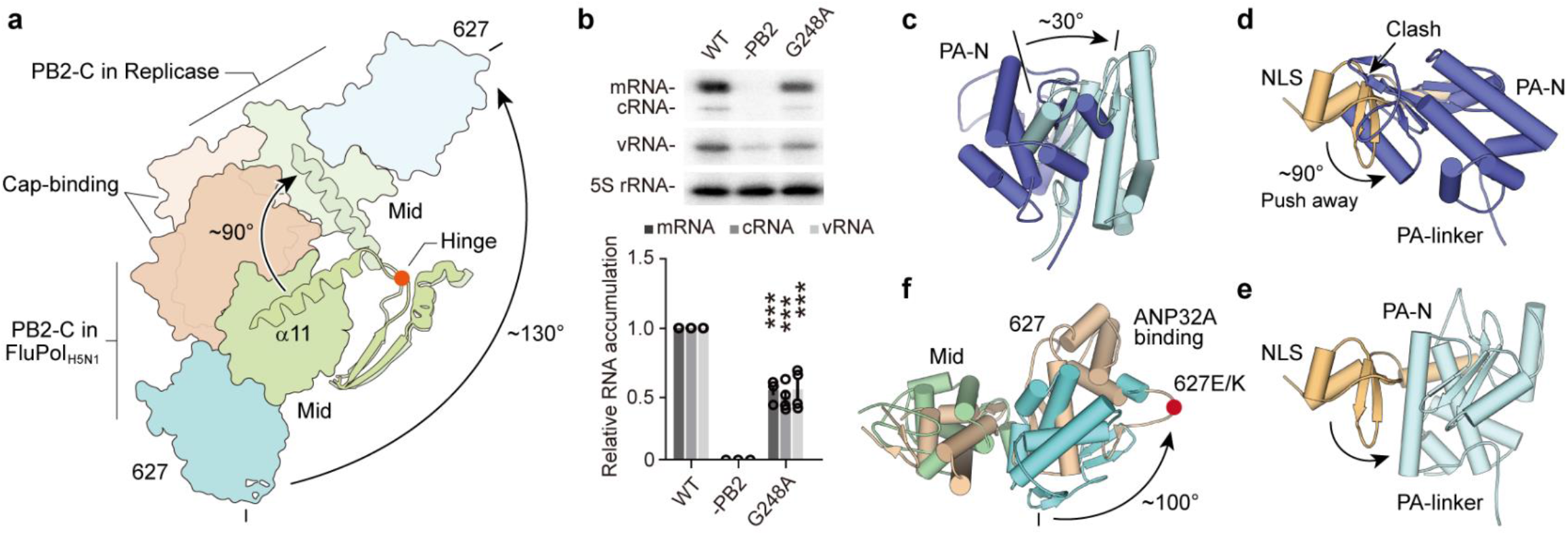
Transition from the resident FluPol_H5N1_ state to the replicase state. **a**. The conformational changes of PB2-C domains between FluPol_H5N1_ and replicase (PDB: 6XZG) are shown. Domains in FluPol_H5N1_ are shown in darker color, while domains in replicase are shown in lighter color. The entire PB2-C domains rotate ∼130°, mediated by the hinge and the PB2 helix α11. **b**. Mutagenesis and quantitation of PB2 Gly248 using a viral RNP reconstitution assay. *n* = 4 independent transfections. Data are shown as mean ± s.e.m. ***, *P* < 0.001. **c**. Superposition of FluPol_H5N1_ (blue) and replicase FluPol (light cyan) reveals position and rotation changes of PA-N domains. **d** and **e**. After rotating the entire PB2-C domains, PB2 NLS domain (shown in gold) translocate ∼90° away from of PA-N domain in avoiding of steric clash between PB2 NLS and PA-N, as shown in panel **e**. **f**. Superposition of FluPol_H5N1_ (cyan) and replicase (wheat) shows that ANP32A binding induces the rotation of the 627 domain to an exposed position, leaving the host-specific residue 627 highly accessible to the solvent.

Collectively, our analysis showed that the structure of FluPol_H5N1_ represents an inactive conformation, unfavorable for either transcription or replication. In agreement with this result, the 3′-vRNA proximal end of our structure rests outside the template entrance of polymerase (Fig. S3) (*13, 22*). More importantly, these structural analyses showed that the FluPol_H5N1_ structure can be easily converted into either a T-state structure or a R-state structure. The FluPol_H5N1_ structure converts to that in T-state after a contraction of PB2-C caused by the interactions between its internal domains (Fig. 1f). In comparison, it converts to that in R-state after a rotation of the entire PB2-C domain (Fig. 2a). Following with the movement of PB2-C, PA-N also rotates accordingly. In other words, these results suggest that there are two successive steps in the transition process from T-state to R-state, a contraction of PB2-C followed by ∼130° rotation of the domains. Our structure, which represents the final stage of the first step, locates on the midway between the T-R transition. Thus, we therefore termed as *resident state*.

### The resident state is required for both transcription and replication

To investigate whether this resident state affects the transcription and replication cycles of FluPol, we analyzed residues that are crucial and specific to stabilize the resident structure. As shown in Fig. 3a, the resident structure is stabilized by interactions between the Cap-binding domain and the Mid domain, as well as the Cap-binding domain and the 627 domain. Specifically, the residue His432 on the helix α17 of the Cap-binding domain interacts with the carboxyl group of Glu517 on η3 of the Mid domain. The residue Arg436 which is adjacent to the residue His432, interacts with the carbonyl group of Glu516 in the Mid domain. In addition, the carbonyl groups of the main-chain residues in 424-loop (Asn422, Ala424 and Asn425) of the Cap-binding domain interact with the residues Lys586 and Arg589 in η4 of the 627 domain by hydrogen bonds (Fig. 3a, 3b).

**Figure 3.**
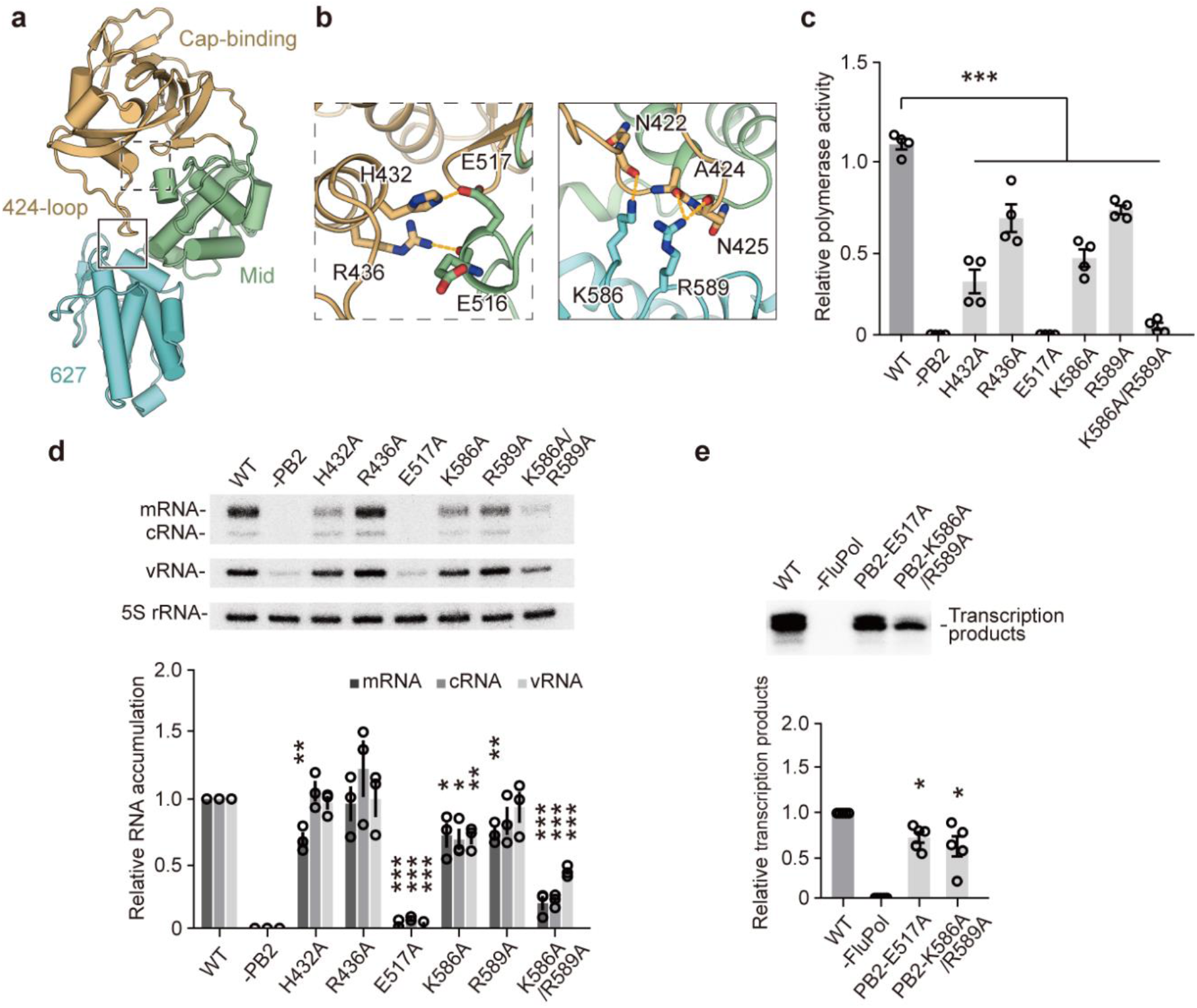
The resident state is critical for both transcription and replication functions. **a**. Structure of the FluPol_H5N1_ PB2-C domain. **b**. Close-up views of the interactions stabilizing the inactive conformation of PB2. Interacting residues are shown in stick representations. **c**. Mutagenesis of stabilizing residues by minigenome luciferase reporter assay. Data are shown as mean ± s.e.m., *n* = 4 biologically independent samples from 4 independent experiments. **d**. Mutagenesis and quantitation of stabilizing residues using a viral RNP reconstitution assay. *N* = 3 independent transfections. Mutations of the residues stabilizing resident state affect the activity of the polymerase complex to produce viral RNAs, with E517A and K586A/R589A double mutant impairing activity most. **e**. Cap-dependent transcription assays. *n* = 5 biologically independent samples. Data are shown as mean ± s.e.m. *, *P* < 0.05; **, *P* < 0.01; ***, *P* < 0.001.

We next generated mutations to destroy the resident structure and tested the polymerase activity of these mutants using standard minigenome reporter assay (*32, 33*). The expression and assembly of mutations, H432A, R436A, K586A, R589A, E517A and K586A/R589A were similar to those of wild type FluPol (Fig. S5). As shown in Fig. 3c, the polymerase activities of H432A, R436A, K586A and R589A single mutations were reduced by approx. 74%, 40%, 60% and 35%, respectively. Notably, the E517A and K586A/R589A double-points mutants displayed abolished polymerase activity (Fig. 3c). Together, these results suggest that the resident state are involved in the functional cycles of FluPol.

To separately study the effects of these mutants on transcription and replication cycles, we evaluated whether the cellular accumulation of positive- and negative-sense viral RNAs were affected in the context of viral RNP, using a RNP reconstitution and primer extension assay (*34, 35*). As shown in Fig. 3d, the transcription cycle was affected by most of the mutants, including H432A, E517A, K586A, R589A and K586A/R589A, indicated by a decrease in viral mRNA accumulation. Importantly, the E517A and K586A reduced both mRNA and v/c RNA accumulations in RNP reconstitution assay. Compared with RNA levels of the K586A mutant, the synthesis activities of different RNAs further decreased in the double-points mutant K586A/R589A (Fig. 3d), consistent with previous reports showing that Lys586 and Arg589 are required for general polymerase activity of influenza virus (*36*). These results suggest that the resident structure is required for both transcription and replication of FluPol.

In cellular context, FluPol synthesizes mRNA using vRNA as a template, and the inhibited accumulation of vRNA may interfere mRNA synthesis. To clarify whether these mutants affect transcription, we next performed the cap-dependent transcription assays *in vitro* using purified recombinant FluPol proteins (Fig. 3e and Fig. S5). Consistent with the cellular data shown in Fig. 3d, our results confirmed that the E517A and K586A/R589A mutants severely decreased the products of transcription of FluPol (Fig. 3e). Together, our results showed that mutations disrupting the resident state impaired the activity of FluPol by severely affecting both transcription and replication of the enzyme. Thus, the resident state locating at the intersection of transcription and replication cycles is indispensable for the functions of FluPol.

### The core region of the FluPol is elastic in transcription and replication cycles

Although the core region of FluPol comprising PA-C, PB1, and PB2-N is considered to be relatively stable both in T-state and R-state (*15, 18, 19*), we found that the core region contracts or enlarges in accordance with the requirement of the transcription and replication processes (Fig. 4), when comparing the structure of the resident state with those in T-state and R-state. FluPol in the resident state presents a closed conformation characterized by a contracted core with a smaller catalytic cavity. To evaluate the elasticity of the core region, we compared the structure of the resident FluPol_H5N1_ with the structures of transcriptase and replicase. As shown in Fig. 4a and Fig. S6, helices α2, α3, α5, α7, α9, α10, α13, α16, α17 and η3 in PB1, as well as helices α9, α16, α19 and α20 in PA move approx. 1 Å to 4 Å, towards the catalytic center of FluPol_H5N1_, compared to the corresponding helices in the transcriptase (*4, 12*) and replicase (*13*). As a result, the diameter of the catalytic cavity of the resident structure decreases to approx. 18 Å, significantly smaller than of approx. 26 Å found in the structures of FluPol at both T-state and R-state (Fig. 4c, 4d). Based on the structure analysis, we conclude that the resident state represents a closed conformation, consisting both contracted core and contracted PB2-C (Fig. 1f). In addition, these results suggest that the whole structure of FluPol is elastic, a process that is dependent on the movement of the secondary structures. To initiate transcription or replication cycle, PB2-C is opened by the binding of Pol II or ANP32A at the initiation step of transcription or replication process, while the core is enlarged upon entering of the 3′-RNA template (*10, 11, 13*). These structural analysis results provides structural evidence for previous hypothesis that the template-product duplex may mechanically enlarges the catalytic cavity (*14*). After a cycle of either transcription or replication, the core region is rendered free of both template and product. Thus, it will return to its closed conformation, ready to initiate another cycle of viral RNA synthesis.

**Figure 4.**
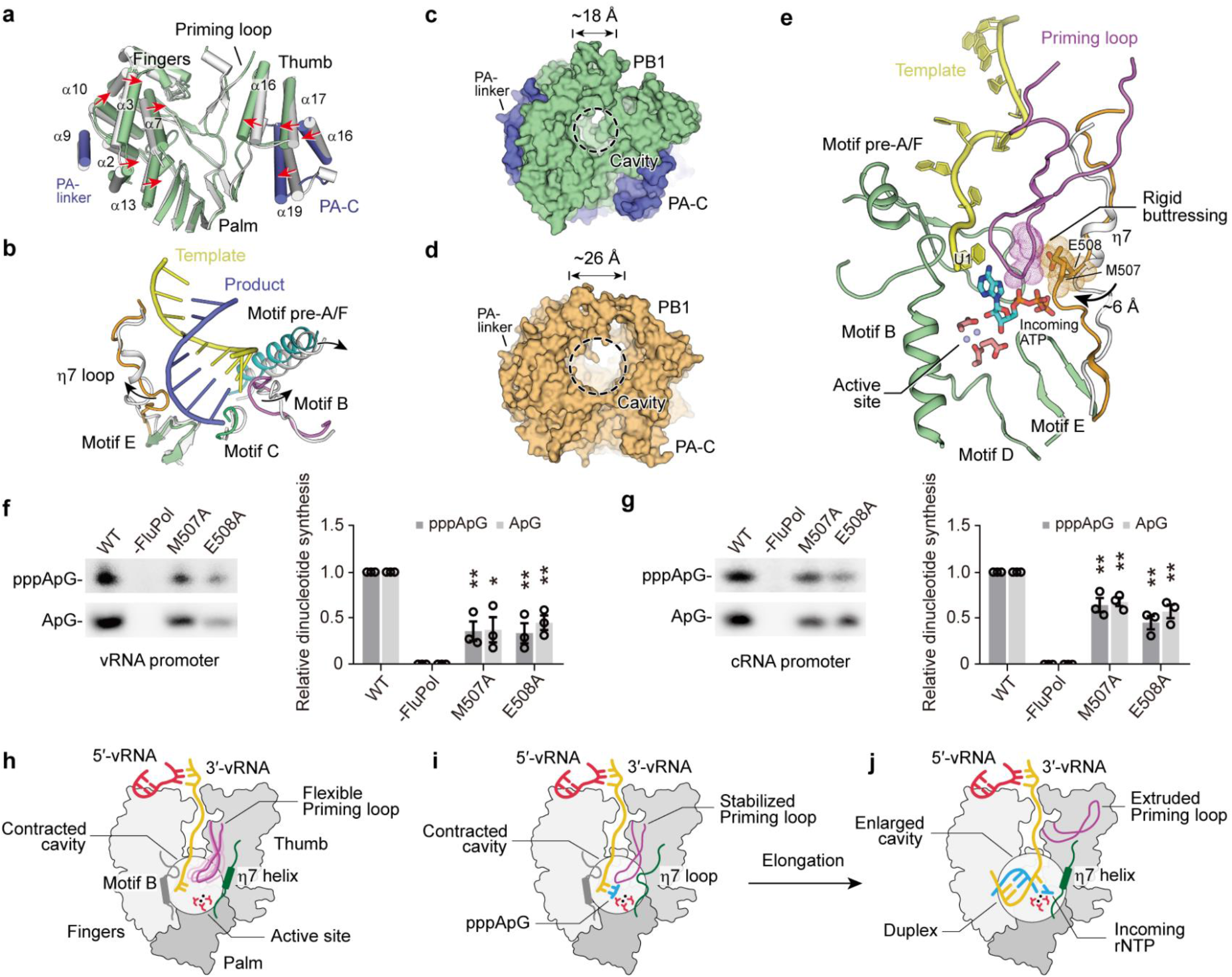
A contracted core of FluPol_H5N1_ is critical for efficient replication initiation. **a**. Structural comparisons of the catalytic cavities in FluPol_H5N1_ (the PA, PB1 and PB2 subunits are colored in blue, green and orange, respectively, as shown in Figure 1**a**) and transcriptase FluPol (PDB: 4WSB, gray). The secondary structures of PA and PB1 in FluPol_H5N1_ are located close to the catalytic sites. **b**. Superposition of FluPol_H5N1_ (with the vRNA duplex) and transcription intermediate (white). PB1 functional motifs are pushed away by the vRNA duplex from the cavity in the presence of template-product duplex. **c** and **d**. Structural comparisons of the cavities in FluPol_H5N1_ (panel **c**) and transcriptase (panel **d**). Structures are shown as surface representations. **e**. The η7 loop stabilizes the priming loop in FluPol_H5N1_ after modeling of template at active site. Met507 and Glu508 on the η7 loop are shown as both stick and dots representations while residues on the priming loop are shown as dots representations. **f**. Terminal pppApG and ApG synthesis on a vRNA promoter of FluPol_H5N1_. **g**. Internal pppApG and ApG synthesis on a cRNA promoter of FluPol_H5N1_. Data are shown as mean ± s.e.m., *n* = 3 biologically independent samples from 3 independent experiments. *, *P* < 0.05; **, *P* < 0.01. **h** to **j**. Model summarizing the roles of the contracted core region and the η7 loop during replication initiation. PB1 functional motifs surrounding the cavity cooperate to ensure efficient replication initiation. However, the flexible priming loop is destabilized the initiating complex (panel **h**). Therefore, the B loop at the N-terminus of motif B stabilizes and corrects the position of the template (*58, 59*) (panel **h, i**), while the η7 loop stabilizes the priming loop by providing a rigid buttressing-like site (panel **i**). Post dinucleotide synthesis, the presence of the η7 loop is incompatible with the presence of the RNA duplex, resulting in η7 loop undergoing a conformational change into the η7 helix upon early elongation (panel **j**). As elongation proceeds, the RNA duplex enlarges the PB1 cavity before causing the prime loop to extrude from the active cavity, as observed previously (*12, 14*).

### The η7 loop in the core region is essential for initiating replication

In addition to the movement of PB2-C and the contractible region, we observed that the priming loop was stabilized by a η7 loop (residues 500-513 in PB1) in the resident state structure (Fig. S5). The η7 loop observed in our structure was folded into an η7 helix in previously reported structures (*4, 12, 22, 31*). The η7 loop (helix) is highly conserved across all influenza viruses. Since stable conformation of priming loop is required to support the template and stabilize initiating complex during *de novo* initiation of replication (*37, 38*), we propose the η7 loop is important for the functions of FluPol. As shown in Fig. 4e, two bulky residues on the bulge of η7 loop, Met507 and Glu508, protrude ∼6 Å towards the priming loop in the cavity and stabilize the priming loop by providing a supporting site.

To test if the stabilizing effect of η7 loop is required for replication initiation, we generated point mutations by substituting Met507 and Glu508 with alanine, and performed a dinucleotide synthesis assay (Fig. 4f, 4g and Fig. S5). We first examined the ability of primer-independent *de novo* replication initiation activity of M507A and E508A mutants using ATP as substrate. When using the vRNA promoter as template, the terminal pppApG synthesis activity of both M507A and E508A were reduced to approx. 35% of that of wild type FluPol (Fig. 4f). Their internal pppApG synthesis activities based on cRNA promoter were reduced to approx. 64% and 45%, respectively (Fig. 4g).

To study if the residues Met507 and Glu508 interact with the triphosphate group of initiating ATP, we performed a dinucleotide synthesis assay using adenosine as substrate. We found that ApG synthesis activity was attenuated to similar levels when using adenosine as substrate. M507A and E508A showed approx. 37% and 45% of the terminal ApG synthesis activity, as well as 67% and 57% of the internal ApG synthesis activity (Fig. 4f, 4g). These results suggested that both Met507 and Glu508 support the priming loop instead of interacting with the triphosphate group of initiating ATP during dinucleotide synthesis. In addition, our results showed that these mutants caused more serious destruction for the terminal ApG synthesis activity, irrespective of ATP or adenosine being used as the substrate (Fig. 4f, 4g). This finding is consistent with previous report showing that the priming loop is more important for terminal initiation on vRNA template than internal initiation on cRNA template during replication (*19, 38*). Together, these results show that η7 loop regulates efficient replication initiation by stabilizing the priming loop.

When we analyzed the residues stabilizing the η7 loop, we found that FluPol regulated the conformations of η7 loop by contracting the core region. The η7 loop was stabilized by residues in helix α16, α17 of PB1 and α18 of PA through both hydrogen bonds and hydrophobic interactions (Fig. S7). Specifically, the residue Asn504 on the η7 loop interacted with Asn537 on helix α16 of PB1 via the hydrogen bond. The residues Leu509 and Phe512 on η7 loop inserted into a hydrophobic pocket surrounded by Val529, Ile530, Leu539 on helix α16, Phe551 on helix α17 of PB1 and Leu586 on helix α18 of PA (Fig. S7). Based on these findings, we hypothesized that the conformational changes of η7 loop was associated with the motion of the core region. The helix α16 and α17 of PB1 moved outwards to accommodate the RNA duplex during the transcription or replication cycles. The shift of helices α16 and α17 disrupted their interactions with the η7 loop, resulting in the conformational changes of η7 loop to η7 helix. In agreement with this result, η7 helix was observed in the structure of FluPol at early stage of elongation (*12*). Taken together, we proposed an allosteric regulation mechanism for the activities of FluPol. To initiate replication, a contracted core with a η7 loop is required to stabilize the priming loop and increase the efficiency for pppApG synthesis (Fig. 4h, 4i). As the elongation proceeds, the RNA duplex would enlarge the PB1 catalytic cavity. Simultaneously, the η7 loop refolds into η7 helix and releases the priming loop. The flexible priming loop is then extruded by the RNA duplex, as shown in previous reports (*12, 14*) (Fig. 4h, 4i).

## Discussion

Here, we propose that a resident state of FluPol is required for the switch between its transcription and replication cycles. We studied the resident state by a cryo-EM structure of FluPol_H5N1_ bound to the vRNA promoter, which represents a closed conformation, characterized by a blocked Cap-binding domain, a rotated PA-N domain, a contracted core region containing a η7 loop as well as a resting 3′-vRNA promoter. Both structural analysis and structure-based mutagenesis results demonstrated that the resident state structure located at the intersection of transcription and replication cycles.

The resident state structure not only links the reported structures in T-state and R-state, but also provides plausible explanations for the observed experimental results. Firstly, our structure suggests that it is sterically favorable for FluPol to initiate transcription from the resident state. The 627 domain in the resident state is accessible for Pol II CTD peptide (transcription stabilizing factor), similar to that in the transcription pre-initiation state (*4, 10*), although the Cap-binding domain in the resident state is blocked. We assume that loading of Pol II CTD peptide to the binding sites of PA-C and 627 domain triggers a conformational change within the 627 domain (*11, 39*), thereby releasing the Cap-binding domain for “cap-snatching” (Movie S1). In line with this hypothesis, Pol II CTD binding to FluPol stabilizes the conformation of FluPol at transcription initiation state (ref) and enhances its viral transcription *in vitro* (*11*). In addition, our results provide structural evidence showing that the core region will be enlarged during transcription, either by Pol II loading or by the entering of 3′-vRNA promoter, as presumed in previous reports (*12, 22, 31*). Furthermore, our results show that the FluPol can spontaneously fold into an inactive state after fulfill a functional cycle. The contracted core region and the protruding η7 loop towards the catalytic cavity in the inactive state has been observed in the partial structures of FluPol_Bat_ and FluPol_B,_ in the transcription product disassociation and recycling states, respectively (*14*) (Fig. S8). Thus, our results provide a mechanism demonstrating how FluPols are recycled after transcription.

Our results show that some major steps are required to initiate replication by the resident FluPol. The major step is a rotation of the entire PB2-C domain by approx. 130° towards the PA-N domain. Subsequently, the PB2 NLS domain pushes the PA-N domain to rotate ∼90° upward to avoid steric clash. From these observations, an interesting question arises, namely what force drives such a large conformational change. Our structural analysis revealed how FluPol_H5N1_ achieve such transformational transition. The rotation of PB2-C is presumably mediated by the PB2 helix α11 (residues 253-272) in the Mid domain and a loop (residues 247-252) on its N-terminus, which serves as a hinge between PB2-C and the core region. The rotation of PB2-C domain was observed in a previously reported replicase structure (*13*). The PA-C helix α14 of the encapsidating FluPol closely contacts this hinge region at PB2 Gly255 (equivalent to PB2 Gly248 in FluPol_H5N1_) of the replicating FluPol. This interaction triggers helix α11 in the Mid domain to undergo a large shift of approx. 90°, thereby initiating the PB2-C domain to rotate by approx.130° anti-clockwise towards the PA-N domain (Movie S2). Consistent with these analyses, our results showed that a single mutation at PB2 Gly248 (G248A) significantly inhibited cRNA and vRNA accumulations in RNP reconstruction and primer extension assays (Fig. 2b), suggesting an important role in FluPol replication. These results are also in line with several previous studies showing that mutations that increased flexibility or weaken interactions with the core region in this hinge and helix α11 (E249G, R251K, D253N, D256G and T271A) significantly enhanced polymerase activity (*40-45*). Subsequently, ANP32A interacts with replicating and encapsidating FluPols with their 627 domains. The interaction of ANP32A_LRR_ triggers a rotation of the 627 domain by approx. 100°, which would destroy the resident conformation by release the 627 domain from the 424-loop and thereby lead to the exposure of 627 domain as that observed in replicase. Thus, we propose that the transition of FluPol from resident state to the R-state is stabilized by the binding of ANP32A and the encapsidating FluPol (replication stabilizing factors) (Fig. 5 and Movie S2).

**Figure 5.**
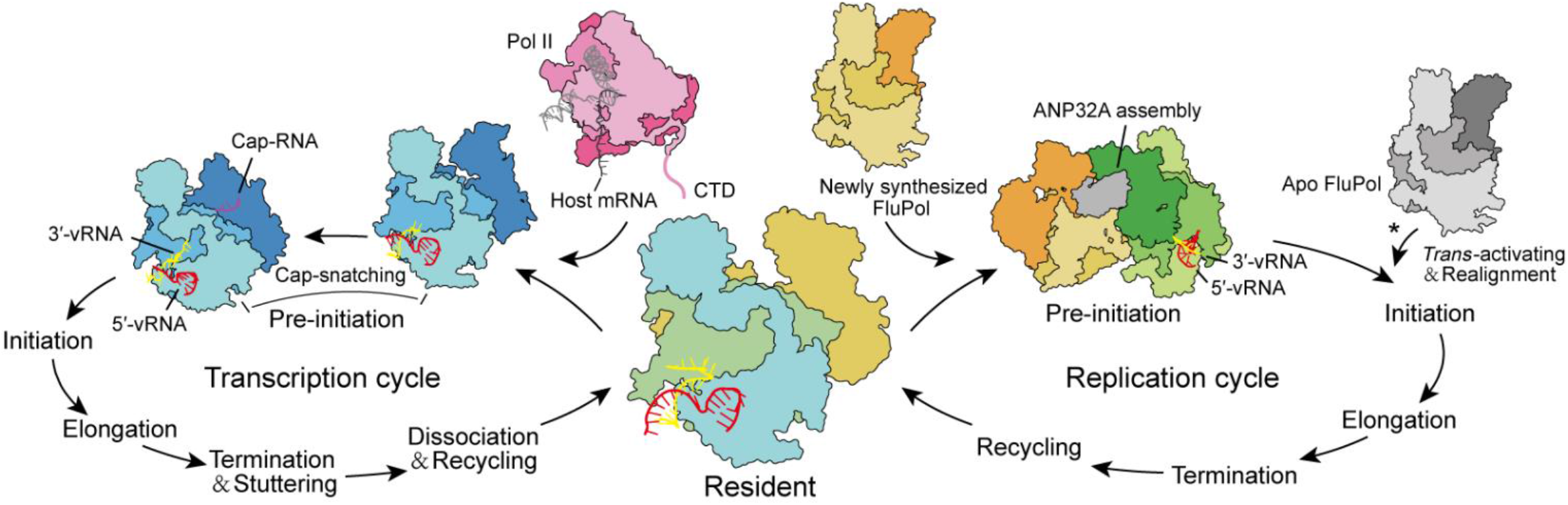
Model of the FluPol resident state-mediated transitions between transcription and replication. FluPol switches between transcription cycle (left part) and replication cycle (right part) via the resident state, a process regulated by host factors. The transition from the resident FluPol to transcriptase is accelerated and stabilized by binding of the Pol II CTD peptide to the polymerase surface. Binding of Pol II CTD to FluPol triggers conformational changes within the 627 domain, thereby releasing the Cap-binding domain and promoting the polymerase to transit to transcription-pre-initiation state. After product dissociation and recycling, FluPol transits back to its intrinsic resident conformation, before starting another functional cycle. As infection proceeds, abundant newly synthesized viral proteins, including FluPol and nucleoproteins, gradually accumulate inside the nucleus. Subsequently, newly synthesized FluPol (called encapsidating FluPol after replicase assembly) enhances the transition from the resident FluPol to replicase, presumably through interactions with the hinge region. Following the rotation of the entire PB2-C and its binding to ANP32A, cellular ANP32A induces the 627 domain to further rotate to an exposed position, rendering the host-specific residue 627 highly accessible, thereby finishing the replicase assembly, and transiting towards the replication-pre-initiation state. Upon completing the replication cycle, FluPol bound with viral RNA returns to its resident conformation, before starting another functional cycle.

In addition, we find that the η7 loop is critical for efficient initiation of replication by stabilizing the priming loop adjacent to the catalytic site. It is known that the priming loop is essential for the functions of polymerase, particularly in replication process (*38, 46*). Increasing evidence suggests that the priming loop should maintain a stable conformation to support the template and stabilize the initiating complex during *de novo* initiation of replication (*37, 38*). However, the reported structures in T-state show that the priming loop is highly flexible, which is extruded out from the core region during transcription elongation (*12, 14*). It was unknown how the priming loop is stabilized in the polymerase, thus hindering the understanding of the replication initiation of FluPol. Motifs equivalent to η7 loop (helix) are also found in the catalytic pocket of polymerases of other known negative-strand RNA viruses (*47-57*). Despite variability of their amino acid sequences, all ‘η7 loops’ (helix) in these polymerases possess bulky residues like phenylalanine, tryptophan and arginine, at the position corresponding to those of Met507 and Glu508 in FluPol (Fig. S9). We thus assume that the mechanism for efficient initiation of FluPol is widely employed by related viruses which also rely on a priming mechanism that initiates replication. It is known that there are some characterized functional motifs residing in a viral RNA polymerase, namely Motif A-H (*4, 47, 51, 54*). Due to the unique functional and structural features, we term η7 as Motif I in the resident state of FluPol.

In summary, our results reveal that FluPol concatenates two completely different initiation processes together by regulating the structural conformation of the resident state under different cellular conditions (Fig. 5). At the early stage of infection, the binding of Pol II CTD activates the PB2-C domains of FluPol and stabilizes its conformation towards transcription initiation state, which drives FluPol transfer to transcription cycle. Once newly synthesized FluPol have been accumulated, some resident FluPols choose the path towards replication initiation stabilized by free Apo FluPol and cellular ANP32A. This provides a reasonable explanation for FluPol to orchestrate the transcription and replication in the life cycle of influenza virus. Since the FluPol is proposed to reside as a low energy state which is apt to be activated in a RNP packaged into a virus, therefore, we propose that this resident state of the FluPol would reflect that constituted in the resting RNP.

## Acknowledgments

We thank staffs in the Center of Cryo-Electron Microscopy, Zhejiang University for their assistance during data collection. We thank Dr. Gang Ji, Dr. Xiaojun Huang, Dr. Boling Zhu, Mr. Longlong Zhang and Mr. Deyin Fan in Center for Biological Imaging (CBI), Institute of Biophysics, Chinese Academy of Science for their assistance during data collection and Ms. Tongxin Niu for her computational assistance. We thank Yinzi Ma and Pengyan Xia for their help during sample screening at the State Key Laboratory of Membrane Biology, Institute of Zoology, Chinese Academy of Sciences. We thank the use of Cryo-EM instruments in the Cryo-EM Facility Center of Southern University of Science and Technology. We thank Dr. HongJie Zhang (Core Facility for Protein Research, Institute of Biophysics, Chinese Academy of Sciences) for assisting us with the detection of radioactivity. This work was supported by National Natural Science Foundation of China (31530015, 31870739, 82071346), “Pearl River Talent Plan” Innovation and Entrepreneurship Team Project of Guangdong Province (2019ZT08Y464), Natural Science Foundation of Guangdong Province, China (2020B1515020035).

## Conflict of interest

The authors declare that there is no conflict of interest that could be perceived as prejudicing the impartiality of the research reported.

## Contributions

H. Li, Y. W., M. L., L. G., Q. W., Z. L., J. Z. and X. Z. performed the experiment. H. Li, Y. W., M. L., H. Liang and Y. L. wrote the manuscript. Y. L. and H. Liang designed the experiment. All authors reviewed the results and approved the final version of the manuscript.

## Materials and Methods

### Cells and plasmids

Sf9 and High Five insect cells that purchased from Invitrogen were used for baculovirus propagation and protein expression. Sf9 cells were maintained in Sf-900 II serum free medium (Gibco) and High Five cells were maintained in ESF 921 Insect Cell Culture Medium (Expression System). Human embryonic kidney 293T cells (HEK-293T) were cultured in DMEM (Gibco) supplemented with 10% fetal bovine serum (Gibco) and with 1% penicillin-streptomycin (Gibco). 293T cells were maintained at 37°C in a 5% CO_2_ atmosphere.

The codon-optimized sequences for the three subunits of avian influenza A/Goose/Guangdong/1/1996 (H5N1) virus polymerases were synthesized (GenScript) and cloned into pFastBac expression plasmid for polymerase expression and structure determination. Plasmids pcDNA-PA, pcDNA-PB1, pcDNA-PB2, pcDNA-NP expressing influenza A/WSN/33 virus proteins, as well as the vRNA-expressing plasmids pPOLI-NA-RT, were kind gifts from professor Dr. Xiaofeng Zheng (Peking University). The firefly luciferase reporter plasmid pYH-Luci and renilla luciferase reporter plasmid pRL_TK were gifts from professor Dr. Wenjun Song (Guangzhou Medical University). The plasmid pcDNA-PB2 was subjected to site-directed PCR mutagenesis to construct plasmids expressing mutant PB2 proteins. Introduction of desired mutations was confirmed by sequencing of the entire PB2 genes.

### Protein expression and purification

The full-length genes encoding PA, PB1 and PB2 (His_6_-fused at the C-terminus) of H5N1 virus were cloned into pFastBac vectors and were further packed into three insect viruses using a Bac-to-bac expression system (Invitrogen). The full-length FluPol_H5N1_ complex was expressed by insect cell line High Five after being infected by PA, PB1 and PB2 expressing-viruses for 60 hours. Insect cells were collected and lysed by sonication with ice-cold lysis buffer containing 40 mM HEPES (pH 7.8), 500 mM NaCl, 20 mM imidazole, 10% v/v glycerol and 0.5 mM phenylmethanesulfonyl-fluoride (PMSF). The insoluble fraction was precipitated by ultracentrifugation (20,000 g) for 30 minutes at 4°C and the supernatant was loaded onto a Ni-NTA superflow affinity column (QIAGEN). The Ni-column was then wash three times and eluted with buffer containing 30 mM HEPES (pH 7.8), 300 mM NaCl, 400 mM imidazole and 10% v/v glycerol. The eluted proteins were concentrated and purified by heparin affinity chromatography (GE Healthcare) using buffer A (30 mM HEPES pH 7.8, 300 mM NaCl, 10% v/v glycerol) and buffer B (30 mM HEPES pH 7.8, 1.2 M NaCl, 10% v/v glycerol). The fractions containing FluPol_H5N1_ complex were verified by SDS-PAGE gel and were further loaded onto size-exclusion chromatography (Superdex-200, GE Healthcare) in 20 mM HEPES (pH 7.8), 300 mM NaCl, 10% v/v glycerol buffer. The fractions corresponding to molecular mass of approx. 500 kD (size of FluPol_H5N1_ dimer) were collected and the concentration was determined by Nanodrop (Thermo Fisher).

### Complex preparation

The oligo nucleotides of viral RNA promoter were produced *in vitro* (*60*). The purified FluPol_H5N1_ complex and the vRNA was mixed at a molar ratio of 1:1.2 and then incubated on ice for 30 minutes. The mixture was further purified using a size-exclusion chromatography on a Superdex 200 Increase column (GE Healthcare) with gel-filtration buffer containing 20 mM HEPES (pH 7.8), 300 mM NaCl, 10% v/v glycerol. The fractions corresponding to the molecular mass of approx. 250 kD (size of vRNA-bound FluPol_H5N1_ monomer) were collected and then transferred onto a Superose 6 Increase column (GE Healthcare) in gel-filtration buffer containing 20 mM HEPES (pH 7.8), 300 mM NaCl. The peak fractions were then collected and centrifuged for 30 minutes at 4°C, followed by sample preparation for cryo-EM.

### Cryo-EM data acquisition

4 μL of 0.1 mg/mL freshly purified FluPol_H5N1_-vRNA complex sample was loaded into glow-discharged (O_2_/Ar, 50 s by Gatan Solarus 950) Quantifoil R1.2/1.3 Cu 300 mesh holey carbon grids and plunge-frozen in liquid nitrogen-cooled liquid ethane, using a Vitrobot Mark IV (FEI) operated at 4°C with a blotting time of 4 s (force setting = 2, humidity = 100%). The grids were firstly checked using a Tecnai 20 microscopy (FEI) operating at 200 kV with a Gatan 626 cryo-Holder. Samples which exhibited good vitreous ice, proper particle concentration and distribution were recycled for further data collection. The checked grids were subsequently transferred to a Titan Krios (FEI) electron microscopy operating at 300 kV equipped with a Gatan K2 Summit direct electron detector. Movie stacks for the core region were automatically collected using SerialEM (*61*) with a preset defocus range from −1.5 μm to −3.5 μm in counting mode at a nominal magnification of 29,000×, with a pixel size of 1.014 Å/pixel. Each micrograph stack was exposed for 6 s with a total of 24 frames. To collect the data of the full-length RdRp, we set the slit width for zero loss peak at 20 eV. The camera was in super-resolution mode and the physical pixel size was 1.04 Å (0.52 Å super-resolution pixel size). Movie stacks for full length were automatically collected using SerialEM (*61*) with a preset defocus range from −1.5 μm to −3.5 μm at a nominal magnification of 22,500×. Each micrograph stack was exposed for 5.4 s with a total of 32 frames. The total dose rate was about 50 e/Å^2^ for each micrograph stack.

### Image processing

The overall strategy of image processing was shown in Supplemental Figure S2. Data were manually checked to exclude contaminations and low-resolution micrographs resulting in a total of 3066 (for core region) and 3328 (for full length) micrographs. The micrographs were motion-corrected using MotionCor2 (*62*) and the defocus values were estimated with Gctf (*63*). All further procedures for data processing were performed in RELION2.1 (*64*). Firstly, 10,000 particles were manually picked without bias and classified into 100 classes. Those representing good classes were selected as references for auto-picking. A total of 779,360 (for core region) and 790,730 (for full length) particles were automatically picked in the whole dataset. After several rounds of reference-free 2D classification, approximately 544,055 (for core region) and 415,348 (for full length) particles were selected and further subjected to global angular searching 3D classification. Further, several particles (∼10,000) were selected from each subset to generate two initial models in RELION (*64*) and were low-pass filtered to 40 Å as references. After several rounds of 3D classifications, a total of 331,874 particles for core region and 238,587 particles for full length with well-defined structural features were selected. For 3D reconstruction of core region, the selected particles were subjected to a round of 3D classification into 6 classes with 35 iterations and 3 classes were further selected (275,433 particles). These particles were re-centered and re-extracted and subjected to another round of 3D classification into 14 classes without alignment. Only 3 classes (261,100 particles, 48.0%) with better quality were finally selected and submitted into RELION_REFINE program which gives rise to a 4.1 Å cryo-EM map. Applying a soft-edge mask and B-factor sharpening generated a final 3.6 Å cryo-EM map based on the gold-standard Fourier shell correlation 0.143 criterion. To reconstruct the 3D structure of the full-length FluPol_H5N1_, the selected particles were subjected to a round of 3D classification into 5 classes with 35 iterations and 4 classes were further selected (174,354 particles). These particles were re-centered and re-extracted and subjected to another round of 3D classification into 16 classes without alignment. A total of 156,666 particles (37.7%) with better quality were finally selected and refined to generate a 4.3 Å cryo-EM map. Applying a soft-edge mask, B-factor sharpening and particle polishing produces a final 3.7 Å cryo-EM map based on the gold-standard Fourier shell correlation 0.143 criterion. Local-resolution of the cryo-EM maps of FluPol_H5N1_ complexes were evaluated using ResMap (*65*) in RELION (*64*).

### Model building and refinement

We combined *de novo* modeling and rigid docking of the known structures to generate atomic model for the complexes. To build the model of the core region, we fitted our previously reported crystal structure of PA-C–PB1-N complex (PDB: 3CM8) (*17*) into the 3.6 Å cryo-EM map using UCSF Chimera (*66*) and manually adjusted in COOT (*67*). The side chains of the residues in most regions could be clearly distinguishable. In the flexible regions, we generated *de novo* modeling for model building based on the cryo-EM map. The model of core region was further refined for several cycles in PHENIX (*68*) till the model fits the map well. For model building of full-length FluPol_H5N1_ complex, we docked the structures of PA-N (PDB: 3EBJ) (*23*) and PB2-C fragment (PDB: 5FMM) (*18*) into the 3.7 Å cryo-EM map using UCSF Chimera and manually adjusted in COOT (*67*), according to their chemical properties. Although the local density of PB2-C is relatively poor, the previously published high-resolution crystal structure of PB2-C of H5N1 virus (*18*) fits the map well after manually adjusted in COOT. The refined model of the core region was further fit into the full-length map by UCSF Chimera. Finally, a total of 2,153 residues and 37 nucleotides were built. Structure refinement was performed using PHENIX.REAL_SPACE_REFINE program in PHENIX (*68*) with chemical restraints including secondary structure and geometry restraints. After several rounds of manually and automatically refinement, the refined structure displayed good quality and evaluated by MolProbity (*69*). The FSC curves were calculated between the models and maps using PHENIX.MTRIAGE (*68*). Cryo-EM data collection, refinement and validation statistics are shown in Table S1. All structural figures were prepared using PyMOL (http://www.pymol.org) and UCSF Chimera (*66*).

### Cap-dependent transcription assay

0.4 μM recombinant polymerase was incubated with 0.5 μM 5’- and 3’-vRNA for 1 h. And then the mixture containing 5 mM MgCl_2_, 1 mM DTT, 2.5 μM capped 16-nucleotide RNAs (5’-m^7^GpppGAAUGCUAUAAUAGC-3’, Trilink), 2 U μL^-1^ RNasin (Promega), 1 mM ATP/GTP/CTP, 1 μM UTP, 0.1 μL [α-^32^P] UTP (3,000 Ci mmol^-1^; Perkin-Elmer) was added and incubated at 30°C for 3 h. The samples were quenched at 95°C for 10 min, following by mixing with an equal volume of loading buffer (80% formamide, 1 mM EDTA, 0.1% bromophenol blue and 0.1% xylene cyan). Then the samples were heated again at 95°C for 5 min and immediately put on ice-water mixture for 3min. Finally, the samples were analyzed on a 20%, 8M-Urea-PAGE and visualized by a phosphor-imaging scanner (Typhoon FLA7000IP). All data were analyzed using Image J and Prism 8 (GraphPad).

### *In vitro* dinucleotide replication initiation assay

To evaluate the ability of wide type and mutant polymerase to synthesize a pppApG dinucleotide *de novo, in vitro* dinucleotide replication initiation assay was performed. Reaction mixtures containing 200 ng purified polymerase, 5 mM MgCl_2_, 1 mM DTT, 2 U/μL RNasin (Promega), 1 μM vRNA or cRNA promoter, 1 mM ATP or Adenosine, and 0.05 μM [α-^32^P] GTP (3,000 Ci/mmol; Perkin-Elmer) were incubated at 30°C for 3 h. Reactions mixture were terminated by incubation at 95°C for 10 min, addition of an equal volume of 80% formamide, 1 mM EDTA, 0.1% bromophenol blue, and 0.1% xylene cyan, and further incubation at 95°C for 3 min. Reaction products were separated by 20% denaturing PAGE containing 7 M urea in TBE buffer and visualized by autoradiography. ^32^P-derived signals were quantified using Image J. Data were analyzed using Prism 8 (GraphPad).

### RNP reconstitution assay and primer extension analysis

Approximately 1×10^6^ HEK-293T cells in 6-well plates were transfected with 1 μg each of pcDNA-PA, pcDNA-PB1, pcDNA-PB2, pcDNA-NP and pPOLI-NA-RT plasmid encoding neuraminidase (NA) vRNA segment using PEI (Polyscience) according to the manufacturer’s instructions. Cells were harvested 36 h post transfection. Total RNA was extracted using TRI Reagent (Sigma-Aldrich) and dissolved in 30 μL of RNase-free water. RNA concentrations were measured using a DS-11 Spectrophotometer (Denovix). The accumulation of viral mRNA, cRNA and vRNA was analyzed by primer extension assay. Briefly, RNA was reverse transcribed with SuperScript III reverse transcriptase (Invitrogen) using ^32^P-labeled primers against positive-sense mRNA and cRNA and negative-sense vRNA, with a primer against cellular 5S rRNA included as a loading control. Products were separated by 8% denaturing PAGE with 7 M urea in TBE buffer. Gels were dried and visualized by phosphorimaging on a Typhoon FLA 7000 scanner (GE Healthcare). RNA signals were quantified using Image J and normalized to the 5S rRNA loading control. Input vRNA signal, estimated from the “-PB2” control, was subtracted from all subsequent lanes. Data were analyzed using Prism 8 (GraphPad).

### Minigenome reporter assay

Influenza polymerase activity was measured by use of a minigenome reporter assay as described previously (*32*). Briefly, 50 ng each of pcDNA-PB1, pcDNA-PB2, pcDNA-PA, and pcDNA-NP plasmids mixed with a firefly luciferase reporter plasmid pYH-Luci (50 ng) and a renilla luciferase internal control plasmid pRL_TK (10 ng), were co-transfected into HEK-293T cells in 96-well plates and incubated at 37°C. At 24 h post transfection, the luciferase activity was measured using a Dual-Glo™ Luciferase Assay System (Promega) according to the manufacturer’s instructions. Firefly and renilla luciferase bioluminescence were measured using a BioTek plate reader. RNP polymerase activity was normalized against pRL_TK activity. Data were analyzed using Prism 8 (GraphPad).

### Western blot analysis

HEK-293T cells grown in 6-well plates were transiently transfected with 1 μg each of pcDNA-PA, pcDNA-PB1 and wild-type or mutant pcDNA-PB2 plasmids using PEI (Polyscience) and Opti-MEM (Gibco). pcDNA3.1 plasmid was used as negative control. At 48 h post transfection, cells were harvested in RIPA lysis buffer (Beyotime) for 30 min on ice. Total protein extracts were clarified by centrifugation at 13,000× g for 15 min at 4°C, and supernatant were mixed with SDS sample buffer and boiled for 10 min. Equal volume proteins from each cell lysate were separated by 10% sodium dodecyl sulphate polyacrylamide gel electrophoresis for western blot. Rabbit anti-PB2 antiserum produced in rabbits was used to blot PB2. A β-actin mouse monoclonal antibody (CST) was used as the internal control. Goat anti-rabbit and goat anti-mouse antibodies conjugated to horseradish peroxidase (HRP) were used as secondary antibodies. The bands were detected using SuperSignal West Pico Chemiluminescent substrate (Thermo) according to the manufacturer’s instructions and scanned using a Tanon 5200 Imaging System (Tanon).

## Data availability

The cryo-EM maps of FluPol_H5N1_ complexes have been deposited in the Electron Microscopy Data Bank under accession number EMD-31239 (core region) and EMD-31240 (full length). The coordinate for the atomic model of FluPol_H5N1_ complex has been deposited in the Protein Data Bank under code number 7EPH.

## Notes

### Competing Interest Statement

The authors have declared no competing interest.

