## Supplementary information for "A Resident State Allows Influenza Polymerase to Smoothly Switch between Transcription and Replication Cycles"

### Supplementary-Figure, Table and Movie

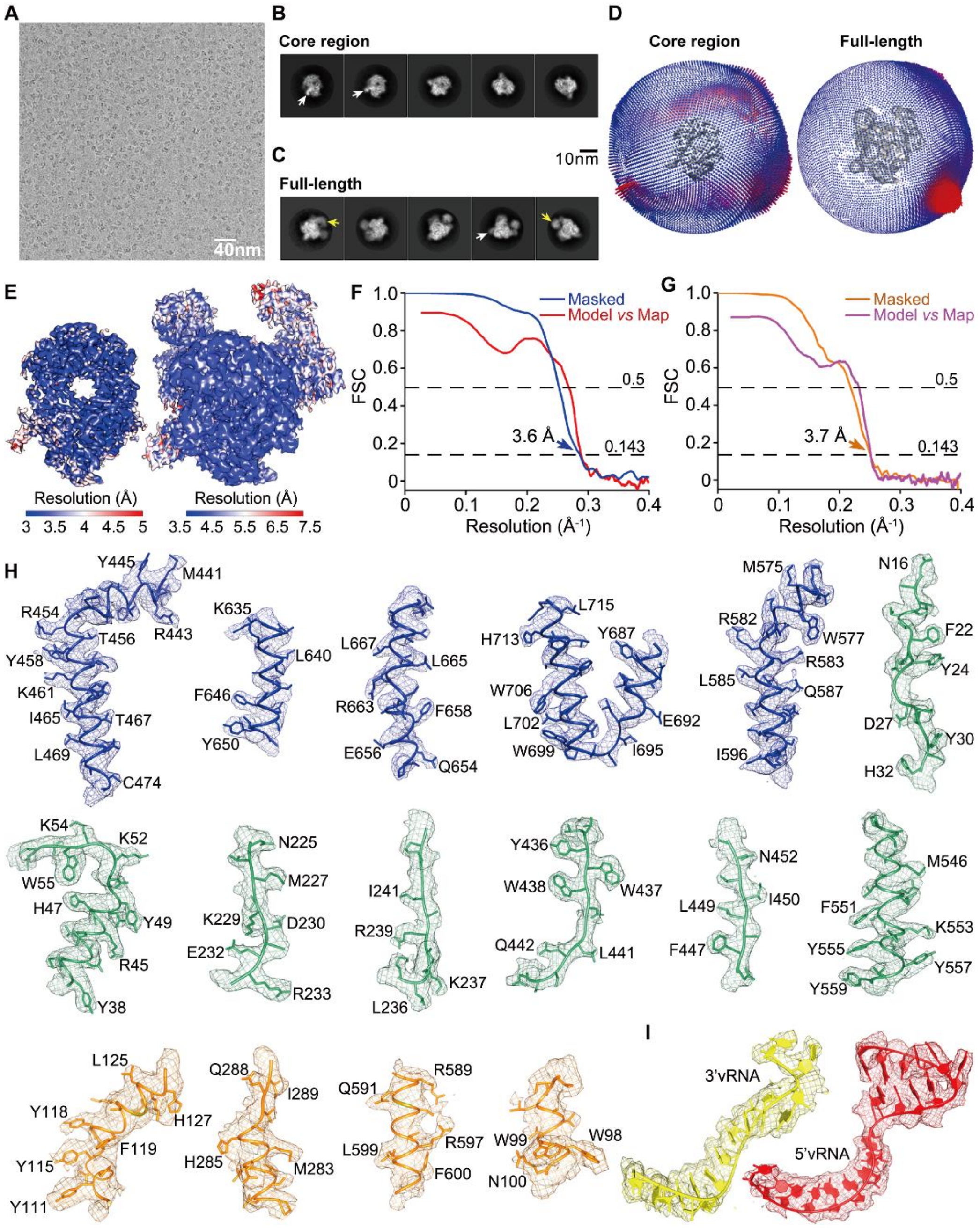

**Figure S1. Cryo-EM analyses of FluPol$_{H5N1}$ bound to vRNA promoter.**

**A**. A representative cryo-EM micrograph of FluPol$_{H5N1}$ bound to vRNA promoter.

**B** and **C**. Representative 2D classifications of FluPol$_{H5N1}$ complex particles. White arrows indicate the vRNA promoter and yellow arrows indicate the flexible domains above the core region.

**D**. Euler angle distributions of FluPol$_{H5N1}$ complex in the final 3D reconstructions.

**E**. Local resolution evaluations of cryo-EM maps of FluPol$_{H5N1}$ complex by ResMap.

**F** and **G**. Gold standard Fourier shell correlation curves for resolution evaluation of core region (**F**) and full-length (**G**) at 0.143 FSC. The FSC curves of the final refined models versus cryo-EM maps at 0.5 FSC are also shown.

**H**. Representative regions of the cryo-EM structure of $FluPol_{H5N1}$ are shown as cartoon representation colored as same as those in Figure 1A. The density maps of motif pre-A (residues 225-243) and motif C (residues 436-453) were shown.

**I**. The density maps of 3′-vRNA and 5′-vRNA.

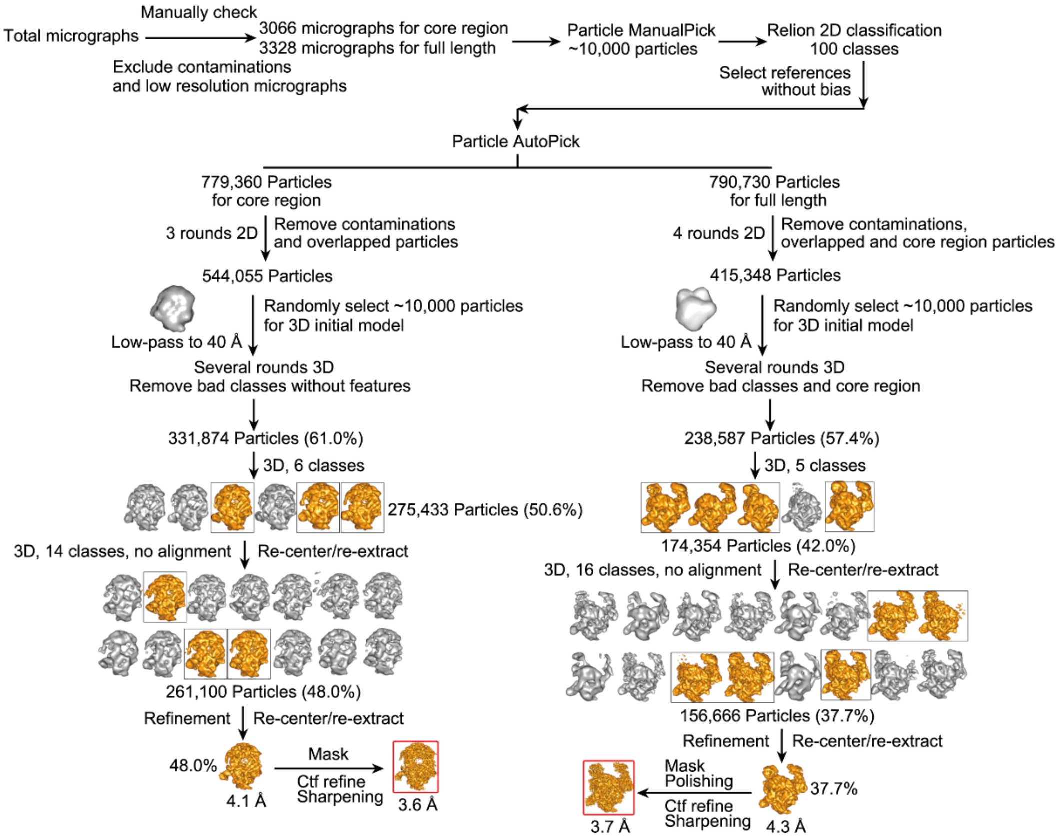

**Figure S2. Data processing strategies for 3D reconstruction of FluPol$_{H5N1}$ complex bound to vRNA promoter.**

The black boxes indicate the selected 3D classes during data processing. The red boxes indicate the final maps.

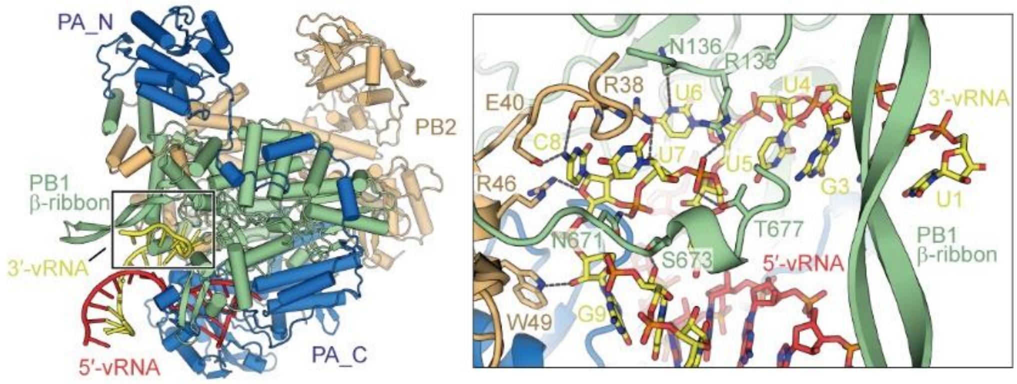

**Figure S3. The 3′-vRNA promoter end resting outside in FluPol$_{H5N1}$.**

The structure of the vRNA promoter in FluPol$_{H5N1}$ complex is shown as cartoon (left) and stick (right) representations. Interacting residues in FluPol$_{H5N1}$ stabilized the 3′-vRNA promoter end are shown as stick representation. For clarity, only polar interactions are shown as black dash lines. Colors are shown as same as those in Figure 1A.

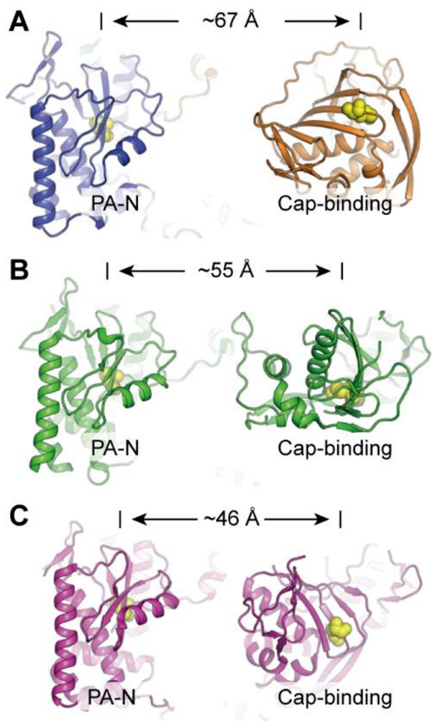

**Figure S4. Distances between PA-N and Cap-binding domains of FluPol in different states.**

The structures of $FluPol_{H5N1}$ (**A**, colored as same as those in Figure 1A), FluPol in transcription-pre-initiation state (**B**, PDB: 4WSB, colored in green) and FluPol in transcription-initiation state (**C**, PDB: 5MSG, colored in magenta) are shown as cartoon representations in top view. Distances between the two active sites (shown as yellow spheres) in PA-N and Cap-binding domains are indicated.

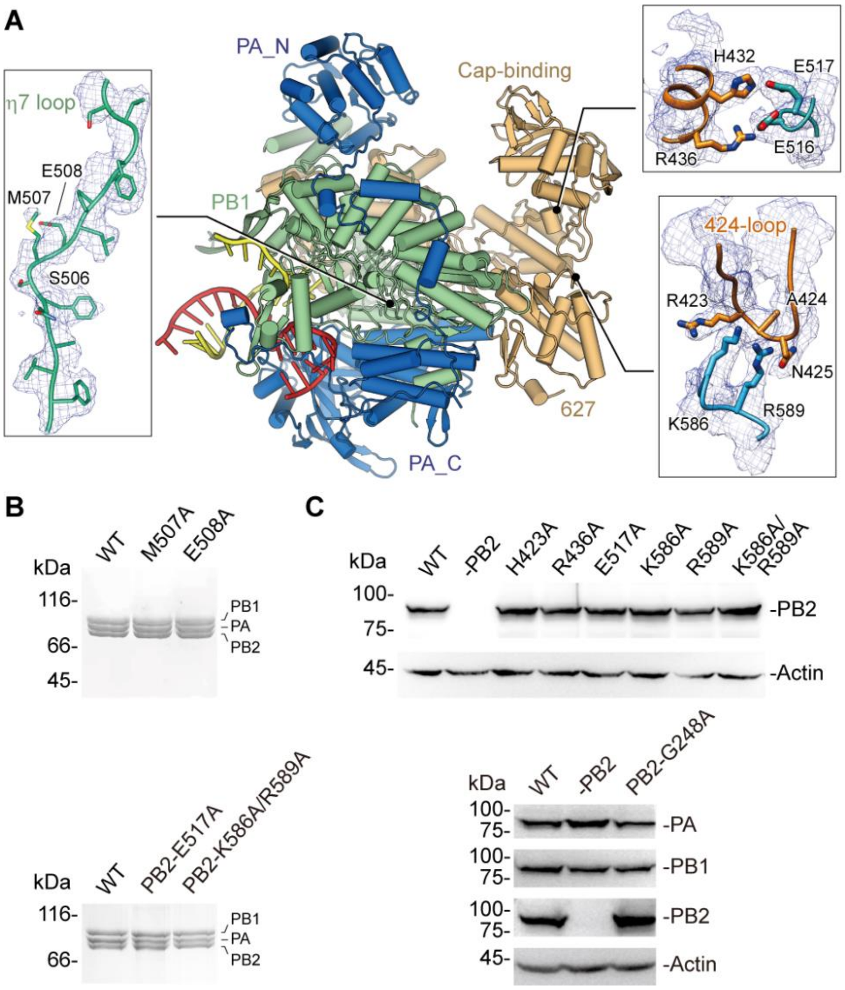

**Figure S5. The density maps and expressions of mutants of FluPol$_{H5N1}$.**

**A**. The density maps of the η7 loop in PB1 (left) and residues stabilizing the inactive conformation of PB2-C FluPol$_{H5N1}$ (right). Colors are shown as same as those in Figure 1A.

**B**. SDS-PAGE analysis of the purifications of mutant proteins (PB1 M507A, E508A and PB2 E517A, K586A/R589A) using the Bac-to-bac expression system.

**C**. Western blotting analysis of the expressions of mutant proteins stabilizing the inactive conformation of PB2-C and the PB2 G248A mutant of FluPol$_{H5N1}$.

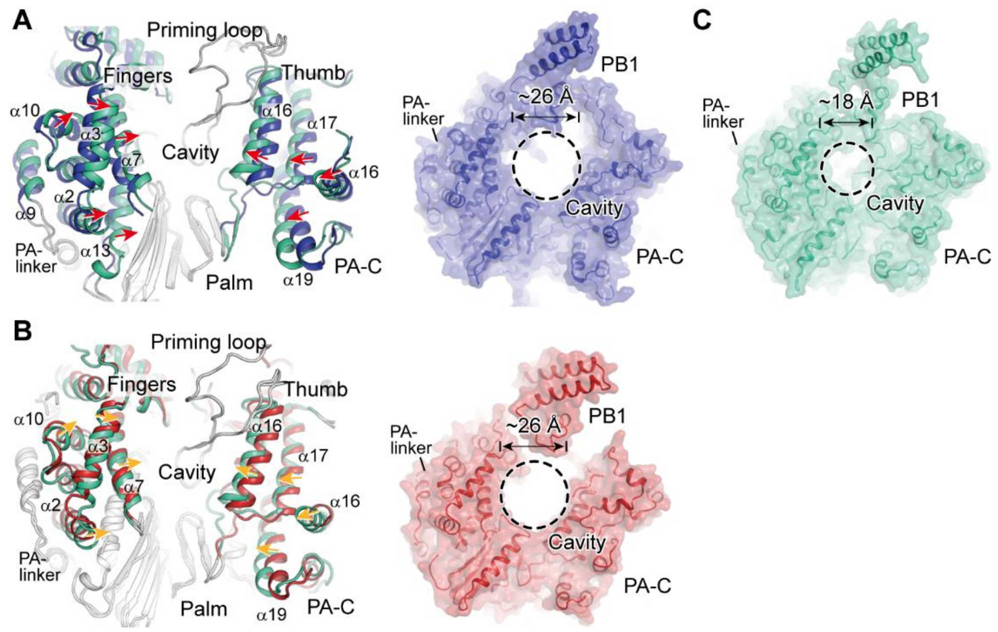

**Figure S6. Structural comparisons reveal a contractible cavity in the resident FluPol$_{H5N1}$.**

**A** and **B**. The secondary structures and the sizes of the catalytic cavities of FluPol$_{H5N1}$ (in green), FluPol in transcription-elongation state (PDB: 6QCT, in blue), FluPol in replicase (PDB: 6XZG, in red).

**C**. Surface representations showing that the cavity of FluPol$_{H5N1}$ is shrunk compared with those in transcriptase and replicase.

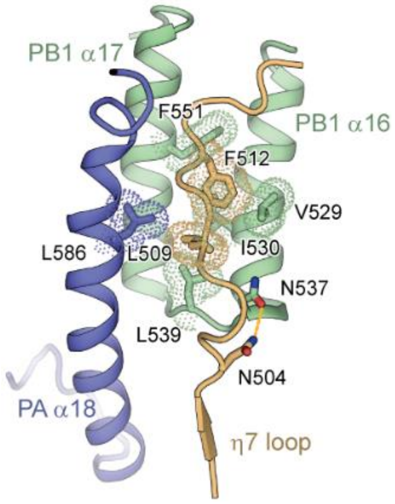

**Figure S7. Interactions stabilized the η7 loop by residues from PA and PB1.**
Structure is shown as cartoon representation and interacting residues are shown as sticks. Hydrophobic interactions are shown as dots while polar interaction is shown as dashed line.

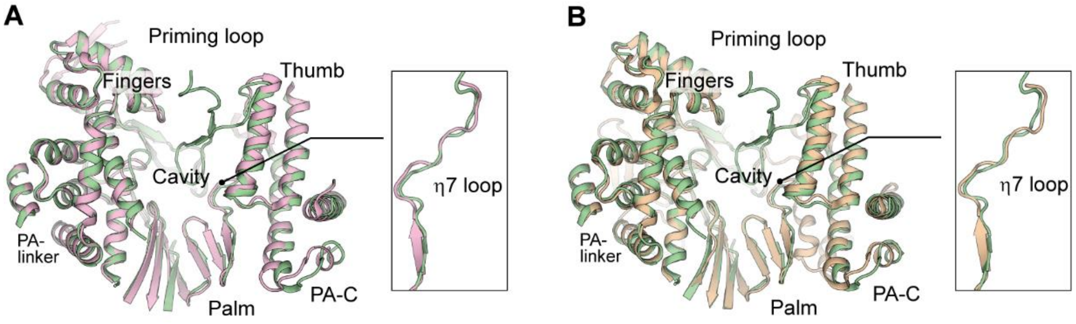

**Figure S8. Structures of core region in product disassociation and recycling states during the end of transcription closely resemble the resident core of FluPol$_{H5N1}$.** The secondary structures of catalytic cavities of FluPol$_{H5N1}$ (in green), FluPol in transcription-product disassociation state (**A**, PDB: 6T0U, in pink), FluPol in transcription-recycling state (**B**, PDB: 6T2C, in wheat). Close-up views show the similar conformation of the η7 loop protruding towards the cavity in these core regions. The priming loop in FluPol$_{H5N1}$ is shown while the priming loops in product disassociation and recycling states are extruded and invisible.

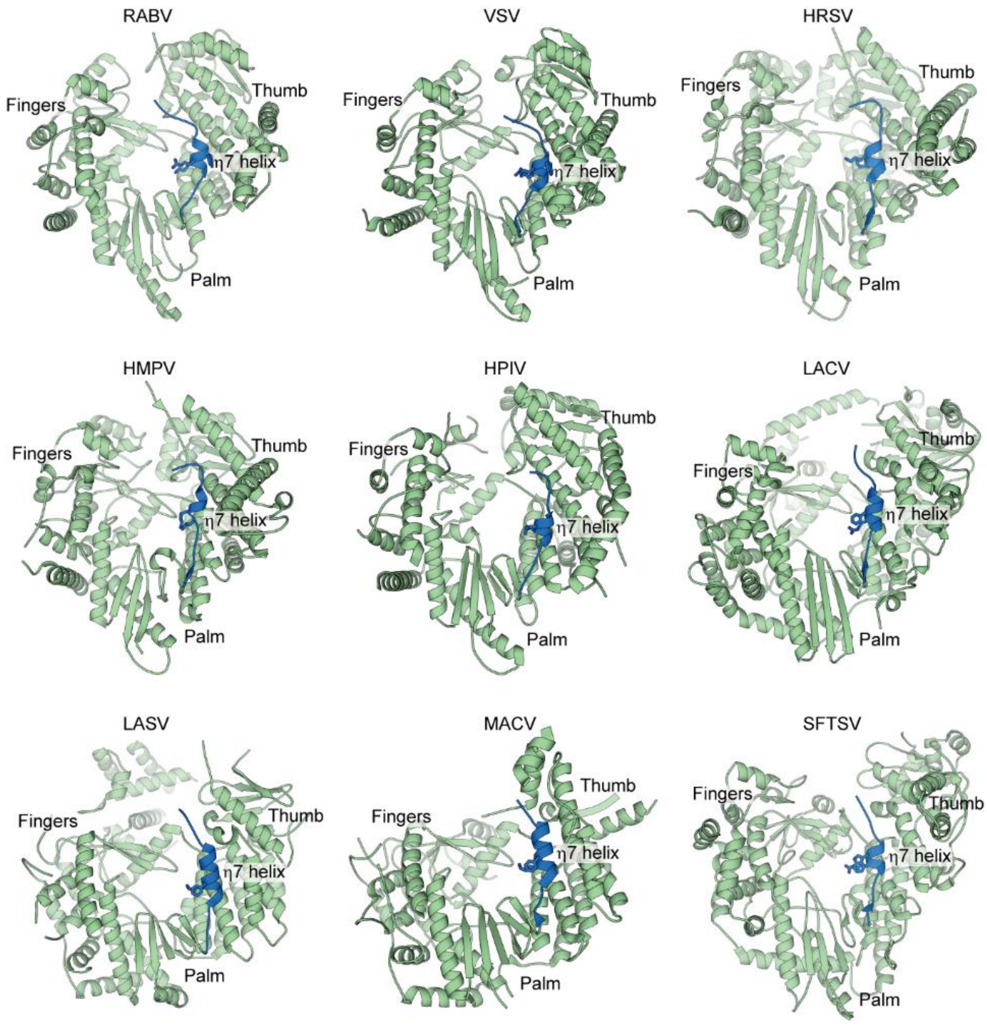

**Figure S9. The η7 helix in the viral polymerase cavity is conserved among different viruses.**

The RdRp regions in RABV (rabies virus, PDB: 6UEB), VSV (vesicular stomatitis virus, PDB: 6U1X), HRSV (human respiratory syncytial virus, PDB: 6PZK), HMPV (human metapneumovirus, PDB: 6U5O), HPIV (human parainfluenza virus, PDB: 6V85), LACV (La Crosse orthobunyavirus, PDB: 5AMQ), LASV (Lassa mammarenavirus, PDB: 6KLC), MACV (Machupo mammarenavirus, PDB: 6KLD) and SFTSV (thrombocytopaenia syndrome virus, PDB: 6Y6K) polymerases are shown as same orientation in cartoon representation (green). The positions of fingers, palm and thumb subdomains are labeled. The homologous structures of η7 helix (in blue) are found in all of these polymerases. Despite of sequence variability, bulky residues like phenylalanine, tryptophan and arginine, are observed at the homologous position of Met507 and Glu508 in influenza polymerase.

**Movie legend**

**Movie S1. Transition from the resident conformation to transcriptase.**

The FluPol is shown as cartoon representation colored as same as those in Figure 1A. The 424-loop in Cap-binding domain and the PB2 helix α11 and hinge are colored in blue and black respectively for clarity. The transition from the resident FluPol to the transcriptase is stabilized by Pol II CTD peptide binding on the polymerase surface. Pol II CTD binding triggers conformational changes of 627 domain, thereby releasing the Cap-binding domain and activating the polymerase for “cap-snatching” towards the transcription-pre-initiation state. Trajectory of intermediate conformations was calculated by UCSF Chimera.

**Movie S2. Transition from the resident conformation to replicase.**

The FluPol is shown as cartoon representation colored as same as those in Figure 1A. The 424-loop in Cap-binding domain and the PB2 helix α11 and hinge are colored in blue and black respectively for clarity. The transition from the resident FluPol to replicase is stabilized by newly synthesized FluPol (called encapsidating FluPol after replicase assembly) through interactions with the hinge region. After the rotation of the whole PB2-C, ANP32A induces the 627 domain to further rotate to an exposed position with host-specific residue 627 being highly accessible, thereby finishing the replicase assembly. Trajectory of intermediate conformations was calculated by UCSF Chimera.

**Table S1 Cryo-EM data collection, refinement and validation statistics**

| | Core region (EMD-31239) | Full-length (EMD-31240) (PDB 7EPH) |
|---|---|---|
| **Data collection and processing** | | |
| Magnification | 29,000× | 22,500× |
| Voltage (kV) | 300 | 300 |
| Electron exposure ($e^{-}$/Å$^2$) | 50 | 50 |
| Defocus range (μm) | 1.5-3.5 | 1.5-3.5 |
| Pixel size (Å) | 1.014 | 1.04 |
| Symmetry imposed | C1 | C1 |
| Initial particle images (no.) | 544,055 | 415,348 |
| Final particle images (no.) | 261,100 | 156,666 |
| Map resolution (Å) | 3.6 | 3.7 |
| FSC threshold | 0.143 | 0.143 |
| Map resolution range (Å) | 3.0-5.0 | 3.0-7.5 |
| **Refinement** | | |
| Initial model used (PDB code) | | N/A |
| Model resolution (Å) | | 3.9 |
| FSC threshold | | 0.5 |
| Map sharpening *B* factor (Å$^2$) | | -120.88 |
| Model composition | | |
| Non-hydrogen atoms | | 18,029 |
| Protein residues | | 2,153 |
| RNA | | 37 |
| *B* factors (Å$^2$) | | |
| Protein | | 146.09 |
| RNA | | 122.25 |
| R.m.s. deviations | | |
| Bond lengths (Å) | | 0.011 |
| Bond angles (°) | | 1.246 |
| Validation | | |
| MolProbity score | | 1.99 |
| Clashscore | | 8.09 |
| Poor rotamers (%) | | 1.36 |
| Ramachandran plot | | |
| Favored (%) | | 93.11 |
| Allowed (%) | | 6.15 |
| Disallowed (%) | | 0.75 |
